# Inhibitory Properties of *Cannabis sativa* Seed Extract on Pancreatic Cancer Cells

**DOI:** 10.1101/2025.07.27.667016

**Authors:** Fatemeh Salah, Mohammad Masoudi, Forouzan Ghasemian Roudsari, Sadrollah Ramezani

**Affiliations:** Department of Biology, Faculty of Science, University of Zanjan, Zanjan, Iran; Department of Biological Sciences, Institute for Advanced Studies in Basic Sciences (IASBS), Zanjan, Iran; Department of Medicinal Plants, Faculty of Geography and Environmental Planning, University of Sistan and Baluchestan, Iran

**Keywords:** Apoptosis, Pancreatic cancer, PANC-1, *Cannabis sativa* seed, Cell cycle

## Abstract

**Background:** Pancreatic Ductal Adenocarcinoma (PDAC) is one of the most lethal cancers and there is a pressing need for development of new therapies. Recently, cannabis has received attentions as a promising plant-based medication for treating a variety of illnesses. The studies investigating the anti-tumor effects of *C. sativa* extract, to date, have used the plant leaves. In this study, we explore the inhibitory properties of *C. sativa* seeds.

**Methods:** An ethanolic extract of *C. sativa* seeds was prepared. GC-MS analysis was performed to identify the compounds in the extract. The cytotoxicity of the extract on PANC-1 cells and HFF cells was assessed using MTT assay. Colony formation and Wound healing assays were used to evaluate the impact of the extract on the ability of PANC-1 cells to form colonies and migrate. Flow cytometry analysis evaluated cell cycle phase of PANC-1 cells after treatment with the extract.

**Results:** The *C. sativa* seed extract had anti-proliferative effects on pancreatic cancer cells PANC-1, while showing no such an effect on normal HFF cells. The extract made its impact on PANC-1 cells by arresting them at G1 phase and increasing their apoptosis. Furthermore it inhibited PANC-1 colony formation and changed the colony combination in favor of paraclones. Migration capacity of PANC-1 cells was also attenuated by the extract.

**Conclusions:** Our results revealed that *C. sativa* seed extract has inhibitory effects on PANC-1 cells by reducing their proliferation, migration and colony formation capacity. It halts the cells in G1 phase and increases their apoptosis.

## 1. Introduction

Among all cancer types, pancreatic cancer is the 12^th^ common cancer worldwide and, due to its poor prognosis, it is the 6^th^ leading cause of cancer-related deaths. The most common type of pancreatic cancer, accounting for more than 90% of diagnosed cases, is Pancreatic Ductal Adenocarcinoma (PDAC, with a 5-year survival rate of less than 5% (Bray et al., 2024). Based on the current data the number of new PDAC cases is increasing by 1% annually (Siegel et al., 2023). PDAC is a particularly aggressive cancer owing to the following factors: rapid growth, high metastatic potential (Le Large et al., 2017; Rhim et al., 2012), poor prognosis and resistance to current treatment options (Binenbaum et al., 2015; Olive et al., 2009). Due to the limited effectiveness of current PDAC treatments, with an average survival rate of only 5-6 months, there is a pressing need to develop new therapies (Bengtsson et al., 2020). Treatment options for PDAC vary depending on the disease progression, and include radiation, chemotherapy, and surgical interventions (Soloff et al., 2018). The surgical approach is an effective treatment for patients with localized tumors in the early stages. However, only a small percentage of patients are diagnosed at an early stage with localized tumors. Consequently, most patients present in advanced stages with metastatic tumors, where treatment options are more limited, and chemotherapy is typically used as the first-line standard treatment for these individuals (Sohn et al., 2000). However, chemotherapy is associated with significant adverse systemic effects. Despite significant advancements in current treatment methods for pancreatic cancer, the impact of these therapies on patient survival remains minimal, and the survival rate has shown little improvement (Kolbeinsson et al., 2023). Accordingly a surge in research dedicated to explore alternative treatment, including ones derived from plant-based sources is ongoing (Jha et al., 2022; Vasques Sr et al., 2024; Zhang et al., 2022).

Cannabis (Cannabis sativa) is an annual plant that belongs to the Cannabaceae family (Kuddus et al., 2013). Recently, cannabis has received a lot of attention as a promising plant-based medication for treating a variety of illnesses (Albarracín & Chaparro, 2018; Mukosi & Motadi, 2023; Shin et al., 2024). Recent studies have demonstrated that active compounds derived from this plant exhibit pharmacological properties, including antimicrobial and antioxidant effects (Baranauskaite et al., 2020; Maayah et al., 2020; Shin et al., 2024; Tang et al., 2009). Typical pharmacological studies have investigated *C. sativa* antimicrobial and analgesic activities (Tang et al., 2009) antidiabetic properties (Baranauskaite et al., 2020; Vasques Sr et al., 2024), and anticancer activity (Bala et al., 2019; Erhabor et al., 2024; Maayah et al., 2020) anti-inflammatory and antidepressant effects (Shin et al., 2024). Furthermore, it is effective against oxidative Stress (Tang et al., 2009) and neuropathic pain (Erhabor et al., 2024). In a study aimed at investigating the cytotoxic effects of aqueous cannabis extracts on cell growth and proliferation, and determining the potential anti-cancer effects of this plant, it was demonstrated that the extract showed growth-inhibitory potential against human breast cancer cells MCF-7 and human prostate cancer cells DU145 cell lines (Erhabor et al., 2024). Additionally, the extract inhibited angiogenesis, nitric oxide production, and oxidative stress in the studied cells (Bala et al., 2019; Erhabor et al., 2024). Many studies have shown the anticancer effects of extracts from different types of cannabis against various tumors (Bachari et al., 2023; Baram et al., 2019; Blal et al., 2023; Cherkasova et al., 2024; Emhemmed et al., 2022; Hosami et al., 2021; Hussein et al., 2014; Motadi et al., 2023; Romano et al., 2014; Rožanc et al., 2021; Seltzer et al., 2020; Zhang et al., 2019). Cannabis can affect the tumor microenvironment through initiating anti-tumorigenic cell signaling, regulating apoptosis and angiogenesis (Erhabor et al., 2024), and adjusting the release of different immunogenic and inflammatory cytokines (Yekhtin et al., 2022; Zaiachuk et al., 2021).

In a recent study, the anti-proliferative and pro-apoptotic effects of two main *C. sativa* cannabinoids in a combination (THC:CBD) were shown on pancreatic cancer xenograft mouse models (Le et al., 2024). In another study, a new method was used to isolate a class of cannabinoids from the crude leaf extract of *Cannabis sativa*, resulting in the isolation of cannabinoid glycosides. The cannabinoid glycosides had improved anti-cancer activities, especially against pancreatic cancer cell lines MiaPaca-2 and Panc-1 (Nalli et al., 2024).

Due to the presence of phenolic compounds in hemp seeds, the antioxidant, antibacterial, and cytotoxic activities of hemp seed extracts have been extensively studied in many research works (Alonso-Esteban et al., 2022; Metouekel et al., 2024). The presence of bioactive compounds such as polyphenols, flavonoids, and cannabinoids in *Cannabis sativa* seeds has been associated with anti-inflammatory, antioxidant, and anticancer properties. Recently, silver nanoparticles biosynthesized with hemp seed extract have been developed and investigated in a lung cancer cell line. The results showed promising outcomes in terms of their potential effectiveness (Yontar & Çevik, 2024). While these findings suggest the effectiveness of *Cannabis sativa* seed extract silver nanoparticles on lung cancer, to date, the raw extract obtained from hemp seeds has not been directly used as a phytochemical agent against cancer cells. Given the aggressive nature of pancreatic cancer and the need for alternative treatment approaches, investigating the anticancer properties of *Cannabis sativa* seed extract against pancreatic cancer cells is of interest.

Previous studies on the anti-cancer properties of *Cannabis sativa* has mainly focused on its leaf and flower, with special attention to the two major secondary metabolites, Δ9-tetrahydrocannabinol (THC) and cannabidiol (CBD). In this study, we explore the anti-cancer properties of *C. sativa* seeds. Seeds from Cannabis sativa have significant nutritional and medicinal benefits with a distinct chemical composition compared to the leaves and flowers (Leonard et al., 2020; Logarušić et al., 2019; Russo, 2007). Unlike marijuana, they contain low Tetrahydrocannabinol (THC levels (<0.1–1%), which ensure minimal psychoactive effects (Citti et al., 2019). They are used as a traditional snack in Iran beside nuts, especially during winter. Here, we investigate the effect of an ethanolic cannabis seed extract on pancreatic cancer cells PANC-1. We seek to know whether *C. sativa* seed extract shows any inhibitory effect on growth, migration and colony formation capacities of PANC-1 cells. Since PANC-1 cell line has shown a higher resistance to chemotherapy drugs in comparison to other pancreatic cancer cell lines MIA-PaCa-2 and BxPC-3 (Fryer et al., 2011) it was chosen for this study.

## 2. Materials and methods

### 2.1 Preparation of hemp seed extract

*C. sativa* seed were used in this study were kindly provided by Dr. S. Ramezani. (University of Sistan and Baluchestan, Zahedan, Iran). To prepare the *C. sativa* seed extract, dry *C. sativa* partially ground seeds (130 g) were processed by maceration with 80% ethanol (600 ml) for 48h at 4°C in the dark, accompanied by persistent shaking and stirring. The extract was filtered through a cellulose filter paper. Ethanol was evaporated using a rotary vacuum evaporator at 80°C. To evaporate remaining solvent, the extract was kept under a laminar flow hood at the Room Temperature for 3 days. The amount of 3 gram of dried extract was achieved and stored in the dark at 4°C.

### 2.2 GC-MS analysis

The GC-MS analysis was carried out using Agilent technologies, GC-MS, equipped with auto injector, under the following condition: A thermo electron system focus gas chromatograph, integrated with a Polaris Q ion trap mass spectrometer, employing an HP-5MS capillary column (50 m x 0.25 mm; 0.25 μm), was utilized for the analysis. The temperature program for the gas chromatograph oven started at 50 °C and was increased by 2 min every minute until it reached 290°C (min). Helium served as the carrier gas (2ml/min), and the injector and ion source temperatures were maintained at 280 ° C and 290° C, respectively, for one minute. The electron impact mode (70 eV) was used to operate the detector, and the m/z 35-550 a.m.u range was used for detection. The NIST Mass Spectral database was utilized to identify the matching compounds after all chromatogram peaks were examined using the Xcalibur® Software. To calculate the retention indexes (RI), a standard alkane solution for GC (C8-C20 in n-hexane) was used. The relative percentage of each component was computed using the GC peak areas without correction factors.

### 2.3 Cell culture

PANC-1 cells were obtained from Iran National Genetic Depository Centre and cultured in high-glucose DMEM medium supplemented with 10% fetal bovine serum (FBS). Cells were cultured at 37°C in a humidified atmosphere containing 5% CO2. HFF cells were obtained from Royan Institute and were cultured in DMEM high-glucose media with 15% FBS.

### 2.4 MTT assay

The cytotoxicity of the *C. sativa* seed extract on cell viability and proliferation of PANC-1 cells was evaluated using MTT assay. PANC-1 cells were seeded into 96-well plates 3×10^3^ cells/well. After 24 h of incubation, the medium was removed, and the cells were treated with different concentrations of *C. sativa* seed extract for 72h. After that, 10μl of MTT solution (5 mg/ml) was added to each well, and the plate was incubated for 3h at 37 °C. The media was discarded and 100μl DMSO was added to each well to dissolve the formazan crystals, and the absorbance was measured at wavelength of 570 nm using a microplate reader.

### 2.5 Crystal violet staining

After 72h of treatment with the extract, the media and cell debris were removed, and the cells were gently washed with phosphate buffered saline (PBS). Cells were fixed with 10% paraformaldehyde at room temperature for 30 min. After washing, 0.02% (w/v) crystal violet solution was added to each well and incubated at room temperature for 5 min. The dye was then removed, and the cells were washed with PBS.

### 2.6 Colony-Forming assay

PANC-1 cells were treated with *C. sativa* seed extract for 48h. Cells were then trypsinized. After carefully pipetting into single cells, both groups (treated and control) were seeded in 6-well plates (200 cells/well). After two weeks, the colonies were fixed with formaldehyde 0.2% (v/v) for 30 min. The plates were stained with 0.05% (w/v) crystal violet for 30 min at RT. The stained colonies were counted and examined to determine the type of colony.

### 2.7 Cell migration

PANC-1 cells were cultured in 6-well plates until 90% confluence and then treated with 1% mitomycin C for 8h at 37 °C to stop cell proliferation. At the center of each well, the cell monolayer was scratched with a pipette tip. Then the culture media was removed, the cells were washed gently twice with PBS to remove dead cells and debris, and the cells were imaged using an inverted microscope at 0h, 24h, 48h and 72h. The ImageJ software (version 1.8.0) was used to examine the scratched surface area. Normalized percentage of migration was determined by following equation:

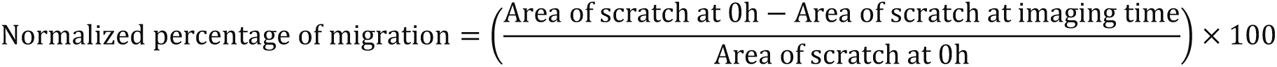

### 2.8 Cell cycle analysis

PANC-1 cells were plated in 60 mm dishes (2×10^5^ cell/dish). After 24 h the cells were treated with *C. sativa* seed extract 1.3 mg/ml for 48h at 37°C. The cells were detached from plates and fixed with cold 70% ethanol at 4°C for overnight. Subsequently, the cells were washed twice with PBS. Next, the cells were simultaneously treated with Rnase A (50 μl of 100 μg/ml) and Propidium Iodide (PI) (200 μl of 50 μg/ml). The cells were then incubated in dark at 37 °C for 1h. Cells were analyzed using a flow cytometer at wavelength of 488 nm and the results were examined using FlowJo (version 10.5.3).

#### Statistical analysis

Statistical analysis of the cell viability assay, colony formation assay and migration assay was performed using either t-test or one-way ANOVA followed by Dunnett’s post hoc test. For colony types and cell cycle assays the chi-square test was used. Stars were representatives of following p-value numbers * (P ≤0.05), ** (0.01< P), *** (P <0.001), **** (P < 0.0001).

## 3. Results

### 3.1 Extract solubility and its effect on PANC-1 Cells viability

We examined solubility of *C. sativa* seed extract in three different solvent, deionized water, PBS and DMSO. Solubility of *C. sativa* seed extract in DMSO was 70 mg/ml and in deionized water was 26.5 mg/ml while it did not dissolve in PBS (Supplementary Information Figure. S1). We then assessed the effects of both DMSO- and deionized water-dissolved extracts on cell viability. There was no difference between the viability of the cells treated with the *C. sativa* seed extract and DMSO as the vehicle (Supplementary Information Figure. S2). However, *C. sativa* seed extract neutralized cytotoxicity effect of DMSO on PANC-1 cells at a high concentration (Supplementary Information Figure. S2). Deionized water-dissolved extract in contrast, showed a significant difference with control (Figure. 1). And based on these results, the extract dissolved in deionized water was used for further experiments. In all the figures presented in the main text, the water-dissolved extract and water (as vehicle) are used for treating the cells. The IC50 value of the *C. sativa* seed extract, against PANC-1 was calculated as 1.3 mg/ml (Figure. 2)

**Figure. 1.**
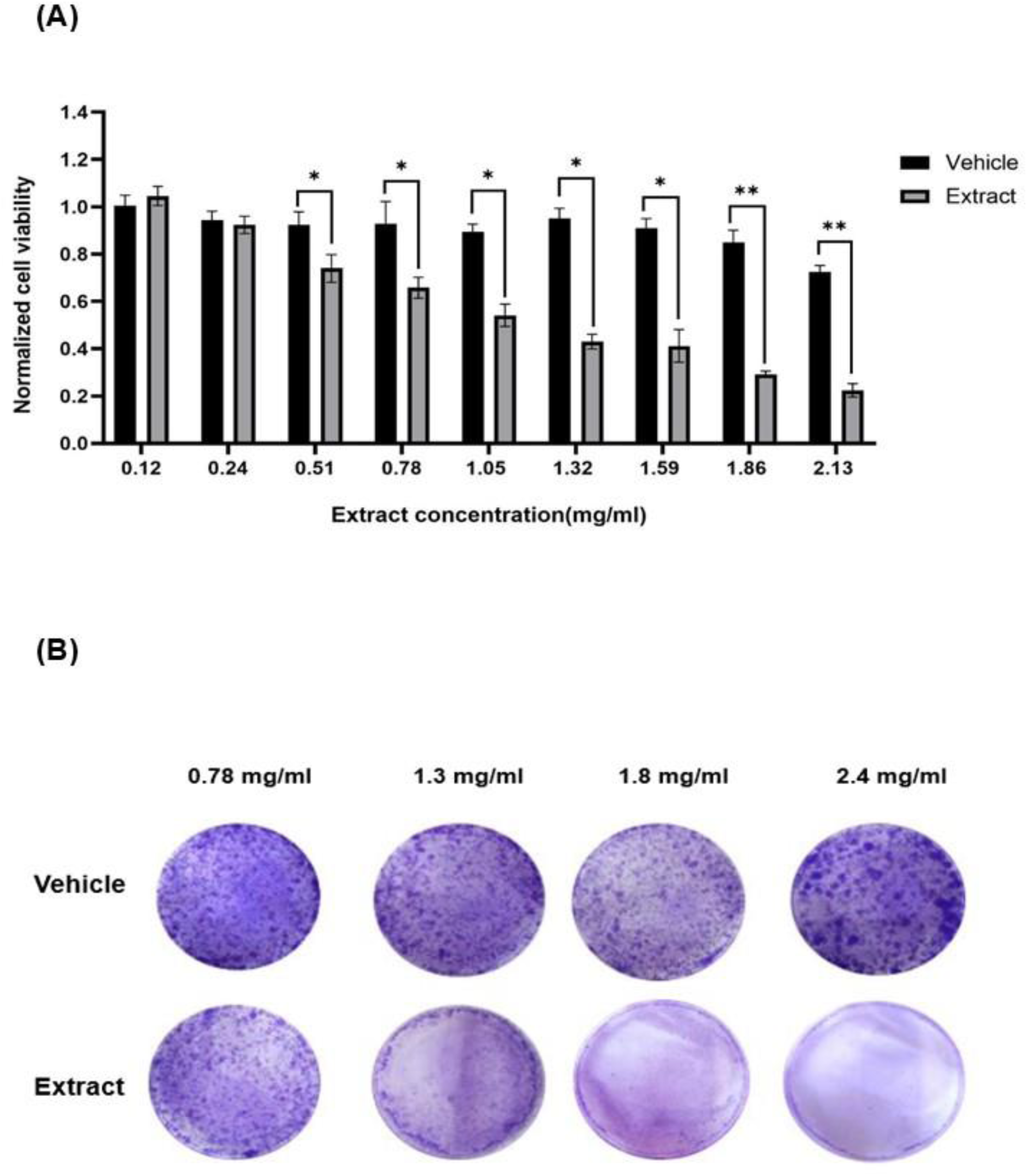
**(A)** The effect of *C. sativa* seed water-dissolved extract on the survival rate of PANC-1 cells were measured using the MTT assay. The cells were exposed to various concentrations of the extract for 72 h. The vehicle is deionized water. Data is representative of three independent experiments. The one-way ANOVA method, followed by Dunnett’s post hoc test was used to analyze the statistical differences between groups. Bars show the mean ± SD of three replicates * (P ≤0.05), ** (0.01< P), *** (P <0.001), **** (P < 0.0001). (**B**) Crystal violet staining of PANC-1 cells treated with different concentrations of the *C. sativa* seed extract for 72h.

**Figure. 2.**
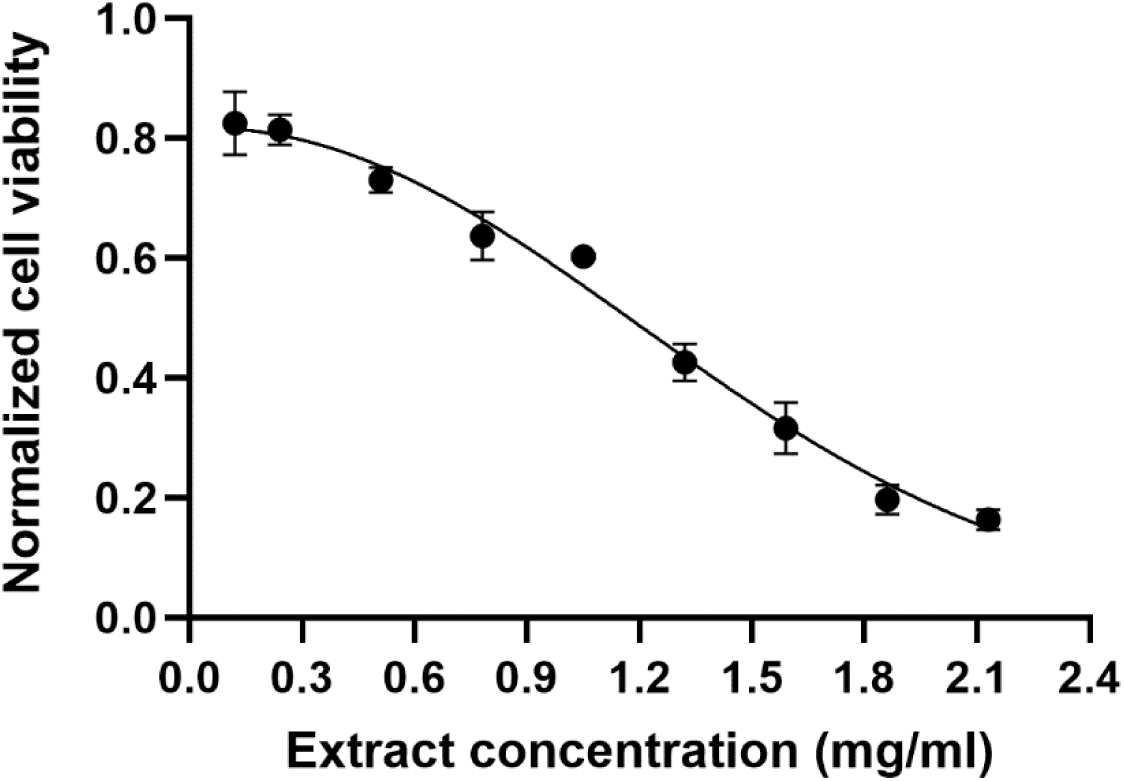
Determining IC50 value of *C. sativa* seed extract on PANC-1 cells. Cells were treated with different concentration of the extract for 72h. Each data point represents the mean ± SD of three replicates. IC50 was calculated as 1.3 mg/ml.

### 3.2 Normal HFF cells viability

Impact of *C. sativa* extract on normal HFF cells were examined by cell viability assay and the extract did not show the growth inhibitory effect on HFF cells at the same concentrations used for treating PANC-1 cells (Figure. 3).

**Figure. 3.**
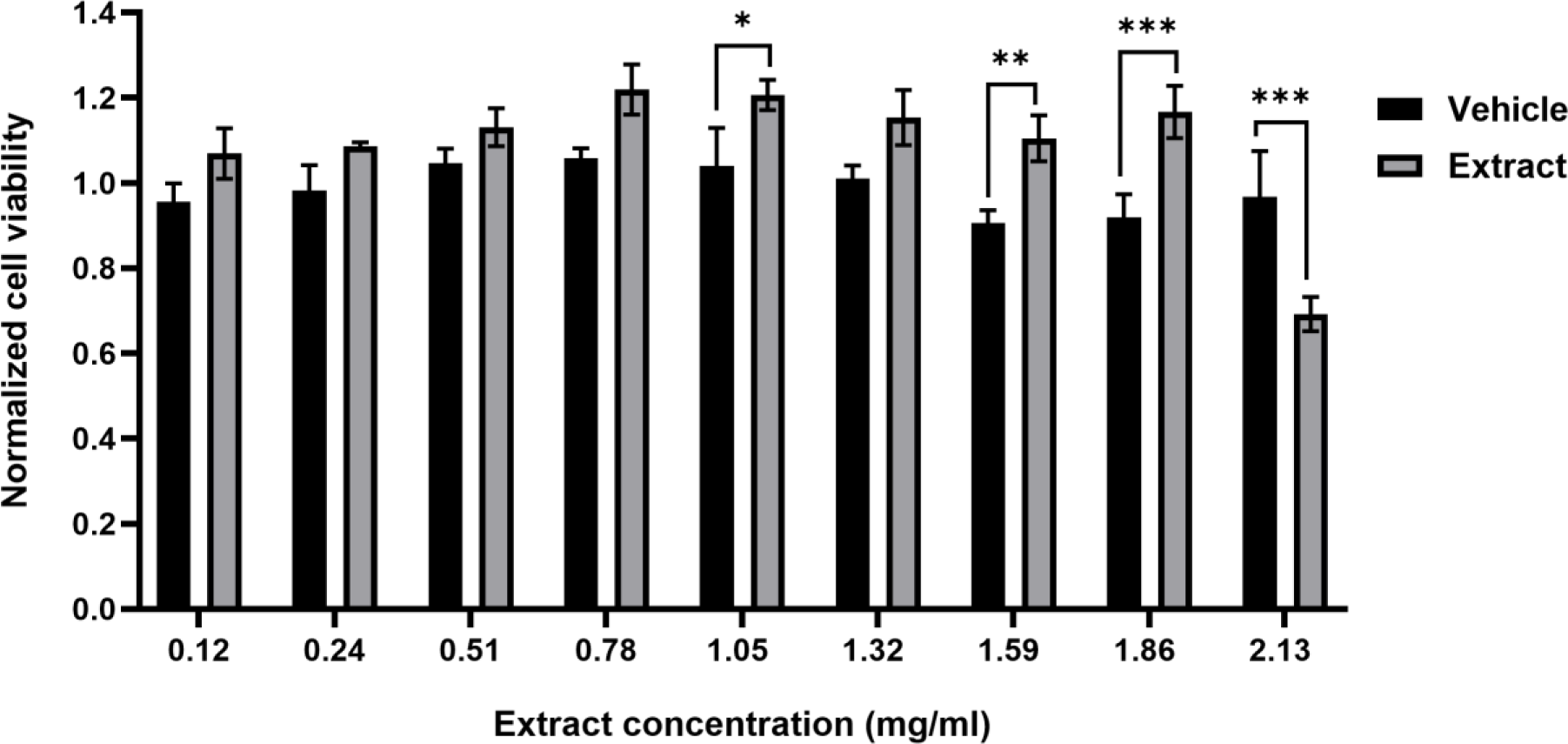
Cell viability assays for normal HFF cells after 72h treatment with water-dissolved *C. sativa* seed extract. The vehicle is deionized water. The one-way ANOVA method, followed by Dunnett’s post hoc test was used to analyze the statistical differences between groups. Data are presented as the mean ± SD of three replicates.

### 3.3 Colony formation

Colony formation assay was used to evaluate impact of the extract on the ability of PANC-1 cells to form colonies. After treatment with *C. sativa* seed extract at concentration of 1 mg/ml the number of colonies decreased in the treated group compared to the control group (Figure. 4). Furthermore, the combination of colony types was changed in favor of paraclones in treated samples (Figure. 5).

**Figure. 4.**
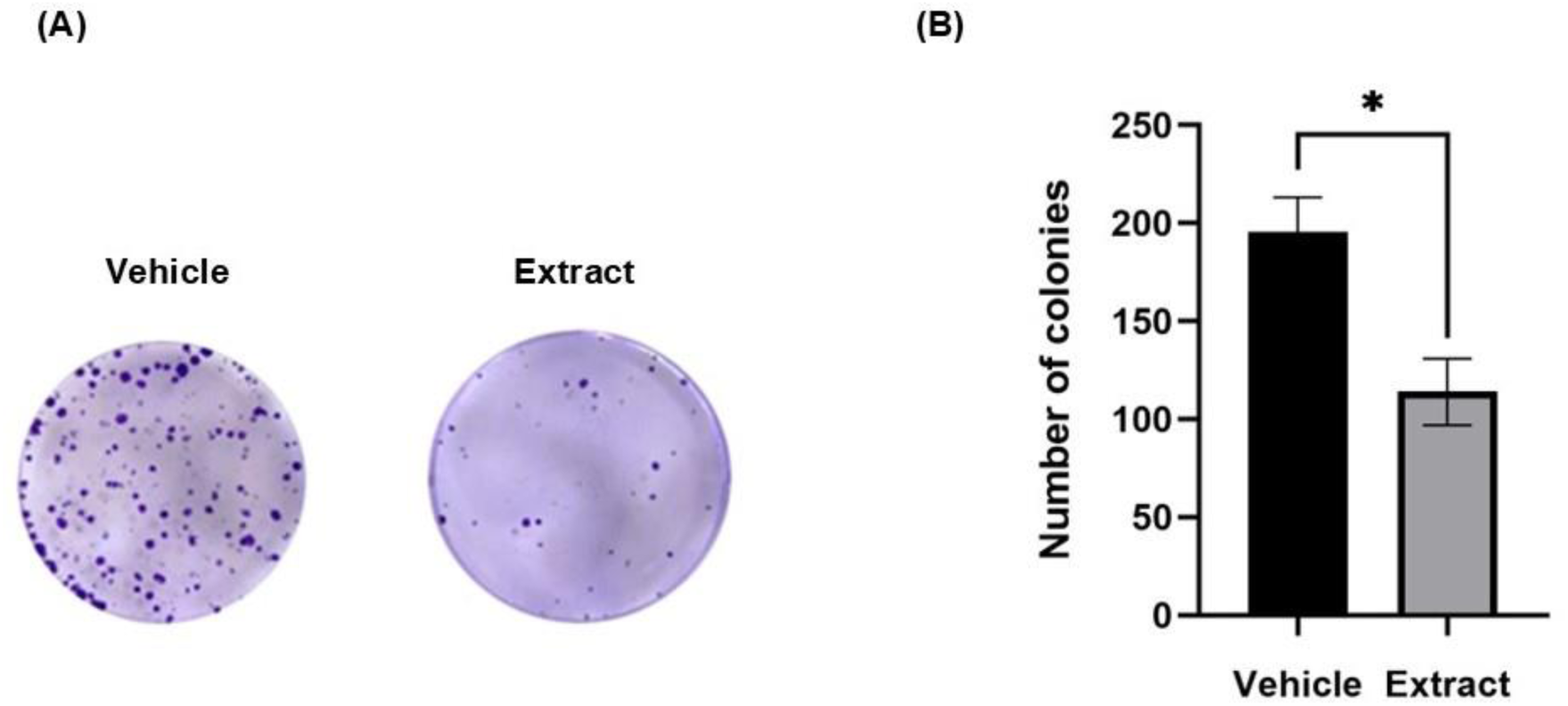
Colony formation of PANC-1 cells treated with water-dissolved *C. sativa* seed extract. The vehicle is deionized water **(A)** Crystal violet staining of the colony formation ability of PANC-1 cells exposed to 1 mg/ml of *C. sativa* seed extract for 48 h. **(B)** Number of PANC-1 cells colony in vehicle- and extract-treated groups. Data are presented as the mean ± SD of three replicates.

**Figure. 5.**
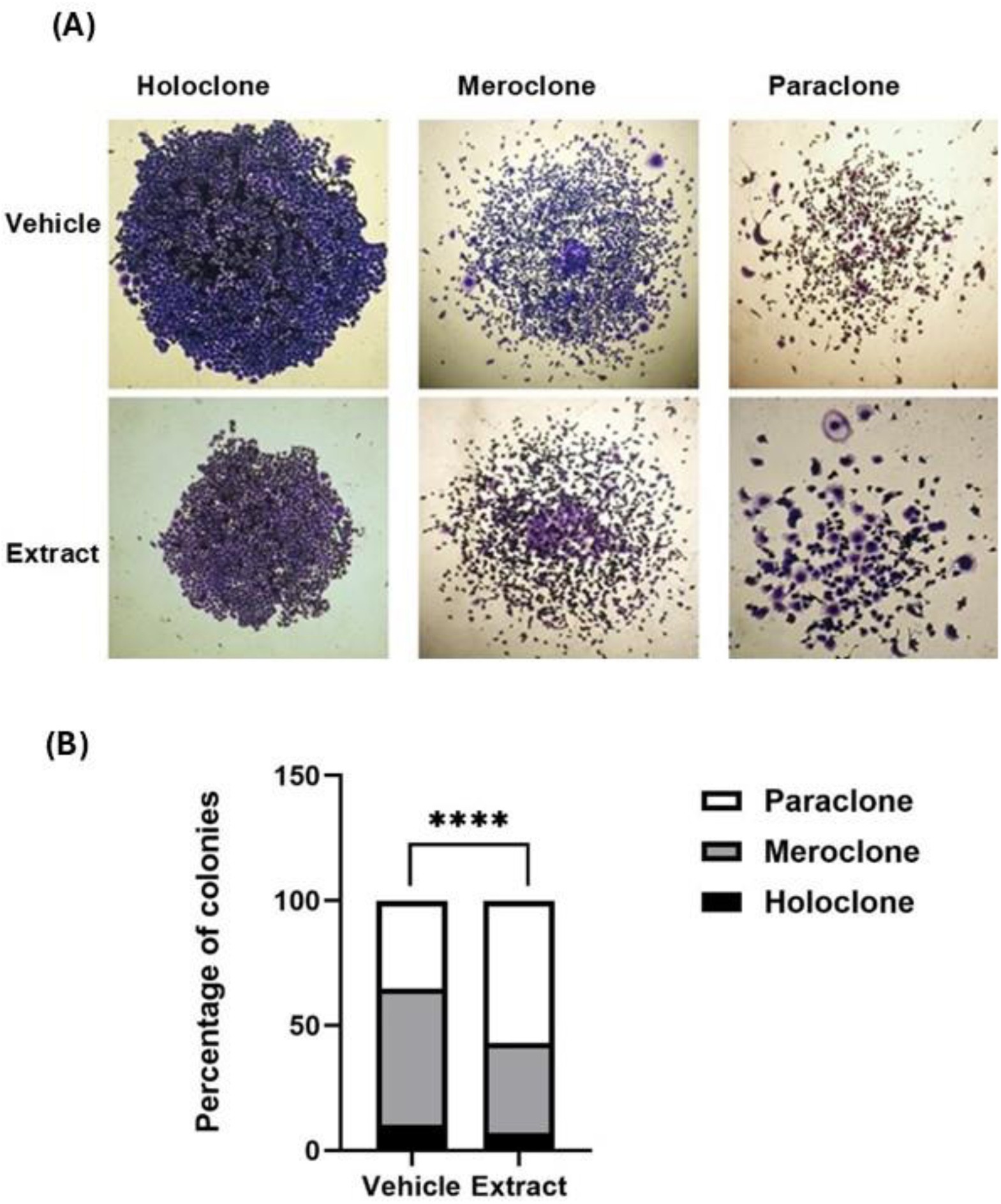
Colony type. **(A)** Morphology of various colony types of PANC-1 cells after treatment with the water-dissolved extract or vehicle (deionized water) for 48h. **(B)** Distribution of various PANC-1 cell colony type after 48h treatment with 1 mg/ml of seed extract.

### 3.4 The effect of C. sativa seed extract on the migration capacity of PANC-1 cells

To evaluate the effect of *C. sativa* seed extract on the migration of PANC-1 cells wound healing assay was performed. Fig.6, shows the results of the scratch assay of PANC-1 cells treated with *C. sativa* seed extract at 0, 24 and 48h. Untreated PANC-1 cells filled up to 96% of the gap space after 48h, whereas extract-treated cells filled up to 68% of the gap space after 48h (Figure. 6).

**Figure. 6.**
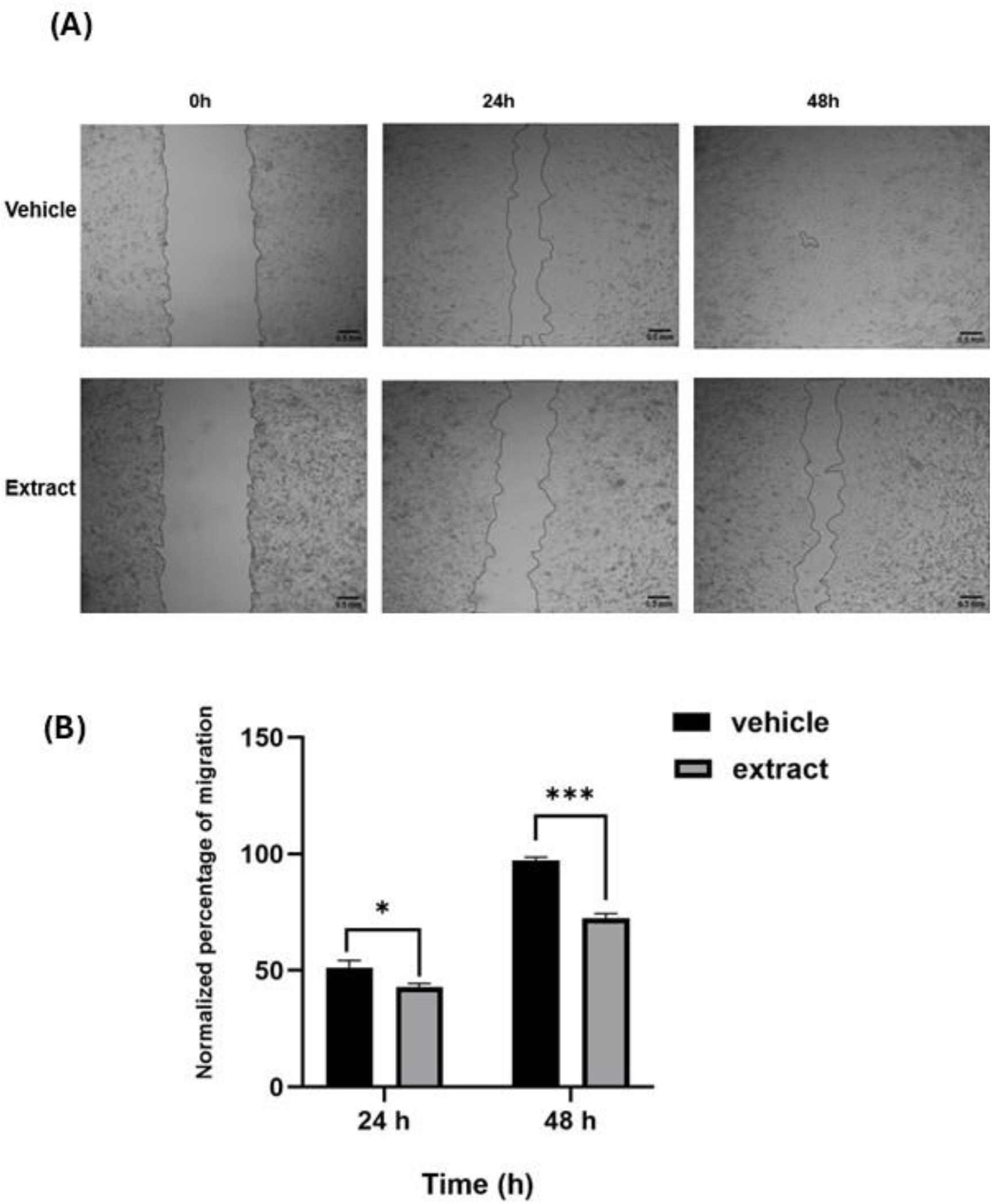
Wound healing assay**. (A**) Microscopic images of wound healing assay at 0, 24 and 48h. PANC-1 cells were treated with 1.5 mg/ml of water-dissolved extract for 48h. The vehicle is deionized water. The cells were treated with mitomycin C to stop their proliferation before scratching. **(B)** Quantitative assessment of the wound area closure. The normalization formula is mentioned in Methods section. The t-test was used to analyze the statistical differences between groups. Data is representative of three independent experiments.

### 3.5 Cell cycle analysis

Analysis of flow cytometry of the cells treated with the C. sativa seed extract in concentration of 1.3 mg/ml showed an increase in the number of PANC-1 cells in the G1 phase compared to untreated cells, along with a noticeable decrease in the percentage of cells in the S and G2/M phases (Figure. 7). These findings indicate that the extract can inhibit the growth of cancer cells via cell cycle arrest in the G1 phase. In addition, the percentage of the cells in sub G1 was increased after treatment with the extract (Figure. 7), indicating an increase of apoptotic cells.

**Figure. 7.**
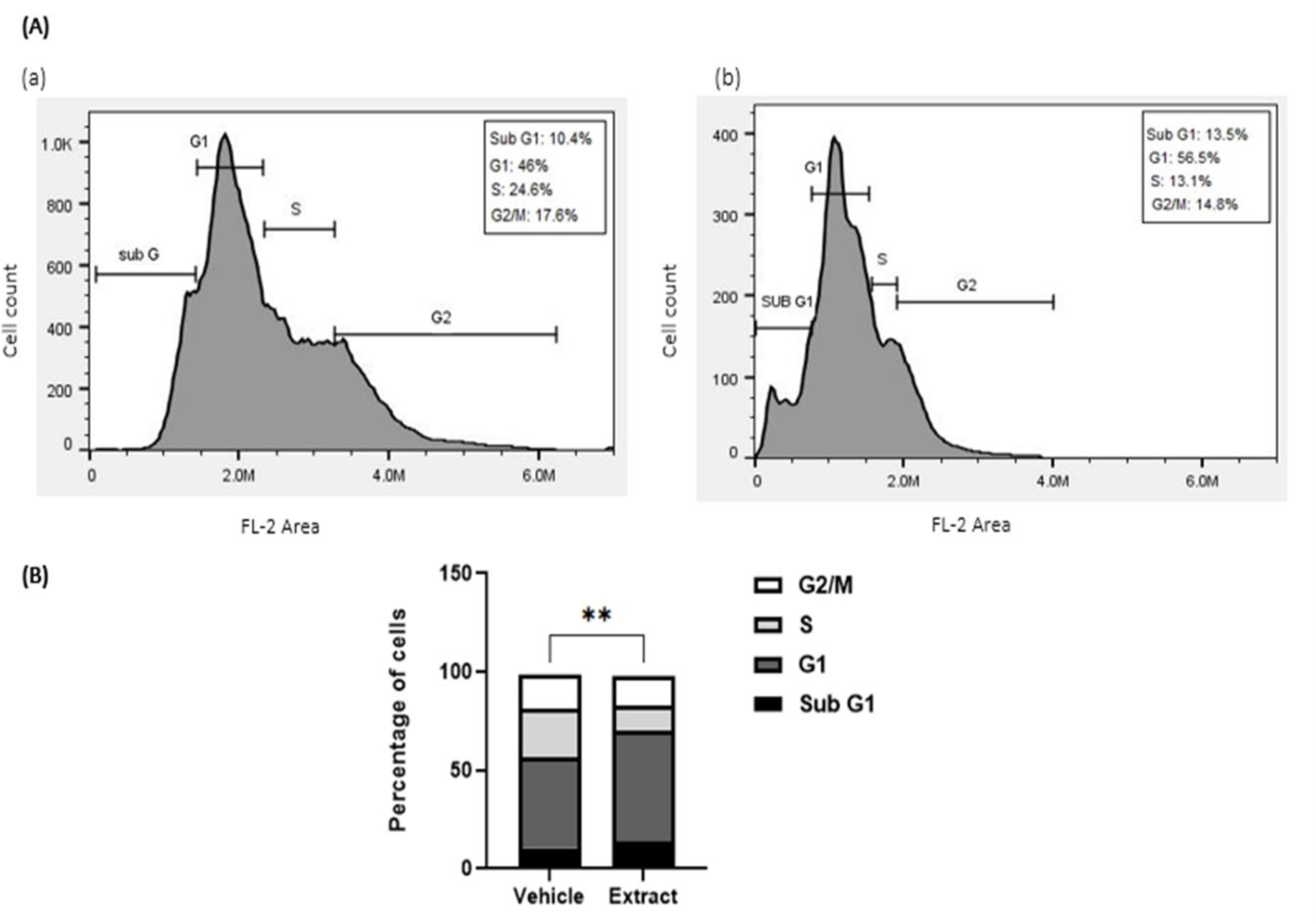
Flow cytometry analysis of PANC-1 cells after treatment with the water-dissolved *C. sativa* seed extract in 1.3 mg/ml concentration for 48 h. The vehicle is deionized water. **(A**) The flow cytometry histograms of cell cycle distribution show the percentage of cells in different cell cycle phases (a) vehicle-treated cells (b) extract-treated cells. **(B)** Plot shows the relative percentage of cells in different cell cycle phases. Data is representative of three independent experiments.

### 3.6 GC-MS Analysis

Quantitative GC-MS data were used to identify the compounds present in the *C. sativa* seed extract, and the amount of each compound was determined through component analysis. A total of 18 compounds have been detected (Table 1). Considering that hemp seeds are oily, a large portion of the compounds identified in the seed extract belong to fatty acids and fatty acid esters (Table 1), which account for 71% of the total identified content in the seed extract. The cannabinoid content identified in hemp seed extract include three types of cannabinoids: The cannabidiol (CBD) 3.83%, Δ9-tetrahydrocannabinol (Δ9-THC) 2.86% and cannabinol (CBN) 1.26%. The total sterol measured in the cannabis seed extract was 6.723%, made up of campesterol, stigmasterol, and γ-sitosterol. The total terpenes identified in the extract were 5.986 %, which consisted of monoterpene β-mono-linolein at a concentration of 5.005 % and phytol at a concentration of 0.981%.

**Table 1.** Identification of the components of the *C. sativa* seed extract in the full spectrum using GC-MS.

| Compound | R.T. (min) | % | Chemical Family |
| --- | --- | --- | --- |
| Desulphosinigrin | 55.14 | 0.08 | Glucosinolate |
| Hexadecanoic acid | 72 | 15.607 | Medium-chain fatty acids |
| Phytol | 78.44 | 0.981 | Terpene |
| cis-9,cis-12-Octadecadienoic acid | 80.19 | 22.754 | unsaturated fatty acid |
| cis-Vaccenic acid | 80.43 | 8.43 | Monounsaturated fatty acids |
| 9,12-Octadecadienoic acid, ethyl ester | 80.75 | 1.458 | Fatty acid esters |
| Octadecanoic acid | 81.17 | 0.398 | Polyunsaturated fatty acid |
| Cannabidiol | 91.72 | 3.835 | Phytocannabinoid |
| 2-mono-Palmitin | 95.26 | 1.867 | Monoglyceride |
| DELTA.9-Tetrahydrocannabinol<br>(Dronabinol) | 95.72 | 2.868 | Phytocannabinoid |
| Diisooctyl phthalate | 95.92 | 1.083 | Fatty acid esters |
| Cannabinol | 98.12 | 1.26 | Phytocannabinoid |
| β-Mono-linolein | 101.9 | 5.005 | Terpene |
| Linolenic acid | 102.16 | 1.167 | Polyunsaturated fatty acid |
| γ-Tocopherol | 113.85 | 1.776 | Tocopherol |
| γ-Sitosterol | 116.94 | 3.865 | Phytosterol |
| Campesterol | 119.39 | 1.94 | Phytosterol |
| Stigmasterol | 120.22 | 0.918 | Phytosterol |

## 4. Discussion

Several studies have shown that the extract of cannabis leaves and flowers can inhibit the proliferation and growth of cancer cells in various cancer cell lines, such as MCF-7, HT-29, PC-3 (Pino et al., 2023), MDA-213 (Bala et al., 2019), HCC1806 (Li et al., 2022), U-87 (Nigro et al., 2020), A549 (Hosami et al., 2021), Caco2 (Rožanc et al., 2021), AsPC-1 (Emhemmed et al., 2022), and HNSCC (Blal et al., 2023).

In this study, we evaluated the cytotoxic effects of *C. sativa* seed extract on the viability of pancreatic cancer PANC-1 cells (Figure 1). The results indicated that the extract dissolved in deionized water significantly reduced the viability of PANC-1 cells in a concentration-dependent manner (Figure. 1). Since the cancer treatment drugs are used systematically and administered drug would reach majority of body normal cells (including normal fibroblast cells) we checked the effect of the extract on normal HFF cells. Many of the side effects of the cancer treatment drugs are due to their effect on normal cells outside their target tissue. Despite the concentration-dependent cytotoxic effect of the extract on PANC-1 cells, cytotoxic effects on normal HFF cells were observed only at the highest concentrations used (Figure. 3). In the current study, we have not investigated the impact of the individual components of the extract; our aim was to examine the effect of the *C. sativa* seed extract as a whole. Nevertheless, we discuss some known effects of the extract’s components for the sake of discussion. Our observation was consistent with a recent study showing that lignanamide-rich fraction of hemp seeds extract exhibited cytotoxic effects on U-87 glioblastoma at a range of concentration, while showing toxicity in normal cells HFF only at the highest concentrations of the extract (Nigro et al., 2020). As shown in previous studies (Metouekel et al., 2024; Raja et al., 2020), the high antioxidant properties of cannabis seed extract might be the reason for the protection of normal cells against the toxic effects of extract.

As evidenced by colony formation assay, the extract caused a significant reduction in the colony formation ability of the PANC-1 cells (Figure. 4). Along with causing a notable decrease in the number of holoclones. Besides a distinct shift from holoclone to paraclone in PANC-1 cells following treatment with 1 mg/ml extract (Figure. 5). Barrandon and Green found that holoclones and meroclones consist of cells that are highly proliferative, immortal, self-renew, and have serial tumorigenic potentials and paraclones are unable to proliferate (Barrandon & Green, 1987). Previous studies, have demonstrated that only holoclones are tumorigenic in vivo (Jeter et al., 2009; Li et al., 2008; Tan et al., 2011) and that holoclones produce larger and faster growing tumors than paraclones (Kalirai et al., 2011). The effect of the extract on reducing the colony formation ability and increasing the percentage of holoclones compared to paraclones suggests that the extract may be considered a potentially useful agent for limiting the colonogenicity of cancer cells.

A lignanamide-rich fraction of *C. sativa* seed extract decreased the colony formation capacity of U-87 cells (Nigro et al., 2020). In our study, treatment of PANC-1 cells with *C. sativa* seed whole extract led to a reduction in the number of colonies, as well as changes in the combination of different colony types (Figure. 4 and Figure. 5).

Cell migration plays a significant role in cancer progression and metastasis. In this study, the ability of cell migration was assessed using the wound healing assay. We found that the extract inhibited the migration of PANC-1 cells after 48h compared to the control. Vaccani et al showed that pure canabidiol has high potential for inhibiting the migration of U87-MG glioma cells (Vaccani et al., 2005). Additionally, Solinas et al (Solinas et al., 2012) demonstrated that this compound had positive effects on reducing endothelial cell migration. Considering that *C. sativa* seeds contain CBD, it might be plausible that the ability of the whole-seed extract to inhibit PANC-1 cell migration is related to this compound.

The results of the investigation into the effect of the extract on cell migration ability were also consistent with recent findings, which indicated that a lignanamide-rich fraction of cannabis seed extract reduced cell migration in the U-87 cell line compared to the control group (Nigro et al., 2020).

We showed that the cannabis seed extract inhibited cell proliferation by inducing cell cycle arrest at the G1 phase and increased cells in the subG1 that is representative of apoptotic cells (Fig. 7). In various cancer cell lines, including A549 lung cells (Hosami et al., 2021), ASPC-1 pancreatic cells (Emhemmed et al., 2022), PC-3 prostate cells (Pino et al., 2023), and head and neck squamous cells (Blal et al., 2023), it has been reported that the extract of cannabis leaves and flowers induced apoptosis. We discovered that the cannabis seed extract influences the apoptosis of pancreatic cancer cells in the PANC-1 line, and the cytotoxic effect of the extract on these cells is related to apoptosis. However, elucidating the detailed mechanisms of cell death caused by *C. sativa* seed extract requires further investigations.

*C. sativa* produces a wide array of secondary metabolites, which serve various biological functions. Many secondary metabolites of this plant have significant medicinal properties. They also exhibit antioxidant (Girgih et al., 2014; Kubiliene et al., 2021), anti-inflammatory (Nigro et al., 2020), and anticancer effects (Koltai & Shalev, 2022; Tomko et al., 2020). Studies on the effects of *C. sativa* extract on various types of cancer (Bachari et al., 2023; Cherkasova et al., 2024; Hosami et al., 2021; Hussein et al., 2014; Koltai & Shalev, 2022; Motadi et al., 2023; Yu et al., 2024) is limited to the leaves of this plant and so far, no study has reported the effects of hemp seeds on any type of cancer specifically pancreatic cancer.

The cannabis seeds are of special interest for us due to their use as a traditional snack in Iran. Given that the compounds and secondary metabolites found in cannabis leaves and seeds are different, they may have various biological effect. We investigated components of the extract we used in this study (Table 1). Cannabis seed contains multiple compounds, including cannabinoids and non-cannabinoid organic compounds, which can play important roles as synergistic and/or entourage compounds (Hanuš & Hod, 2020).

In this study, we analyzed the compounds of the extract that led to the isolation of 18 compounds (Table 1). Although, the aim of our study was to use the whole ethanolic extract of *C. sativa* seeds and we don’t attribute our findings directly to any individual component of the extract, we review some of the known effects of the extract components here.

Hexadecanoic acid is a straight-chain, sixteen-carbon, saturated long-chain fatty acid. In a study conducted by Harada et al, it was shown that the compound N-hexadecanoic acid can selectively inhibit the enzyme DNA topoisomerase-I. Therefore, it can prevent the proliferation of human fibroblast cells (Harada et al., 2002). Based on this finding, Ravi and Krishnan proposed in molecular docking research that N-hexadecanoic acid’s lethal impact results from its interaction with DNA topoisomerase-I, thereby inhibiting cell proliferation (Ravi & Krishnan, 2017). In contrast, Bharath et al. recently demonstrated that hexadecanoic acid induces apoptotic cell death by increasing ROS levels, which causes arrest in the G1 phase (Bharath et al., 2021).

Linolenic acid is mostly obtained from plants, including nuts and seeds. Linolenic acid have anti-inflammatory, antioxidant, and anticancer effects (Yan et al., 2024). Additionally, in combination with anti-tumor drugs, it can enhance treatment efficacy and reduce the side effects of certain medications (Deshpande et al., 2019; İstifli et al., 2019).

One of the most prevalent fatty acid is octadecanoic acid. This fatty acid is present as a glycerol ester in most animal and plant lipids (Beare-Rogers et al., 2001). Farong Yu et al, investigated the cytotoxic and antitumor activity of octadecanoic acid in human gastric (SGC-7901), hepatocellular carcinoma (BEL-7402), and leukemia (HL-60) cells. This compound, in addition to inducing apoptosis and causing cell cycle arrest in the G1 phase, can also lead to necrosis and structural and functional damage to the tumor cell membranes, leading to cell death (Yu et al., 2008).

Previous research has indicated that phytosterols may augment the effectiveness of conventional cancer therapies when utilized in combination treatment regimens. The chemopreventive potential of phytosterols has garnered significant interest within the field of oncology research, with ongoing investigations exploring their applications across various cancer types (Alvarez-Sala et al., 2019; Jesch et al., 2009; Llaverias et al., 2013; Lopez-Garcia et al., 2017; Woyengo et al., 2009).

γ-Sitosterol is a sterol that is considered an isomer of C-24 β-Sitosterol (Thompson et al., 1963). Total sterols content in roots was 0.037%-0.085% and in stem was 0.037%-0.082% (Jin et al., 2021) while here we report a total sterol of 6.7% for the seeds (Table 1).

γ-Sitosterol acts against cancer by inhibiting cell growth, arresting the cell cycle, and inducing apoptotic pathways (Sundarraj et al., 2012). Another study showed that it exerts cytotoxic effects on cell lines by reducing c-myc expression and inducing apoptosis (Endrini et al., 2014).

An increasing number of studies have highlighted the advantages of cannabinoids in cancer treatment, particularly regarding their roles in inducing cell death, inhibiting cell proliferation, and preventing metastasis in various human cancer models, both in vitro and in vivo (Chakravarti et al., 2014; Velasco et al., 2016). Most research conducted on the effects of cannabinoids on cancer focuses on two cannabinoids, THC and CBD (Armstrong et al., 2015; Baram et al., 2019; Drozd et al., 2022; García-Morales et al., 2020; Jeong et al., 2019; Kosgodage et al., 2018; Ligresti et al., 2006; Nabissi et al., 2016; Ramer et al., 2014). THC is the main compound responsible for the psychoactive effects, and it has been measured in inflorescences (10–12%), leaves (1–2%), stems (0.1–0.3%), and roots (<0.03%) (Jin et al., 2020). These amounts vary depending on the plants chemovar, preparation methods, and techniques used for measuring the compounds. However, in general, the THC percentage in the inflorescences is reported higher than that in the other parts. The THC content evaluated in the present study was 2.86%, which was significantly lower than that of inflorescences. CBD is the second most common active compound isolated from cannabis and has potential health benefits, such as anti-inflammatory (Atalay et al., 2019; Wang et al., 2022), anticancer (Massi et al., 2013; Morelli et al., 2014), anxiolytic effects (Zuardi et al., 2017) and neuroprotective properties (Andre et al., 2010). The CBD content was reported 24.9 % and 11.2% in flowers and leaves, respectively, while 0.345% for the inflorescences (Hourfane et al., 2023). In the seed extract examined in this study, the CBD content was 3.835%. The components found in our GC-Ms analysis were similar to the ones previously reported for *C. sativa* seed (Metouekel et al., 2024).

As discussed above, among the compounds identified in the extract, several have been described as molecules with anticancer effects. However, since the goal of our study was to use the whole *C. sativa* seed extract—a combination of these compounds—the mixture may have contributed to the observed inhibitory effects on PANC-1 cells. Identifying the components, or the mixture of components, involved in the observed inhibitory effects of the *C. sativa* seed extract on PANC-1 cells will require further studies. The result of our study, altogether, suggest that the *C. sativa* seed extract might be a candidate for combination therapy alongside known treatments of pancreatic cancer.

## 5. Conclusion

In this study, we demonstrated for the first time that a *C. sativa* seed ethanolic extract has anti-proliferative effects on pancreatic cancer cells PANC-1, while showing no such an effect on normal HFF cells. The extract made its impact on PANC-1 cells by arresting them at G1 phase and increasing their apoptosis. In addition, it inhibited PANC-1 colony formation and changed the colony combination toward paraclones. Migration capacity of PANC-1 cells was also attenuated by the extract. These results suggest inhibitory effects of *C. sativa* seed extract on pancreatic cancer cell line PANC-1.

