## Supplemental Information for "Inhibitory Properties of *Cannabis sativa* Seed Extract on Pancreatic Cancer Cells"

**Supplementary Information (SI)**

**Preparation of the extract solution**

To dissolve the dry extract, three solvents were selected: Deionized water, DMSO, and PBS. Different weights of the extracts were dissolved in 200 µL of each solvent. It was found that the extract did not dissolve in PBS at any of the concentrations, as shown in supplementary Figure 1, while deionized water showed the best solubility at 26.5 mg/ml and the extract showed the highest solubility in DMSO at concentrations of 70 mg/ml. The solubility was defined as dissolving all the extract used in the solvent while leaving no solid sediment left. Based on these primary results, Deionized water and DMSO were selected as solvents for the extract. The dissolution was performed by vortexing and the dissolved extract was filtered through o.22 µm filter and used in following assays.

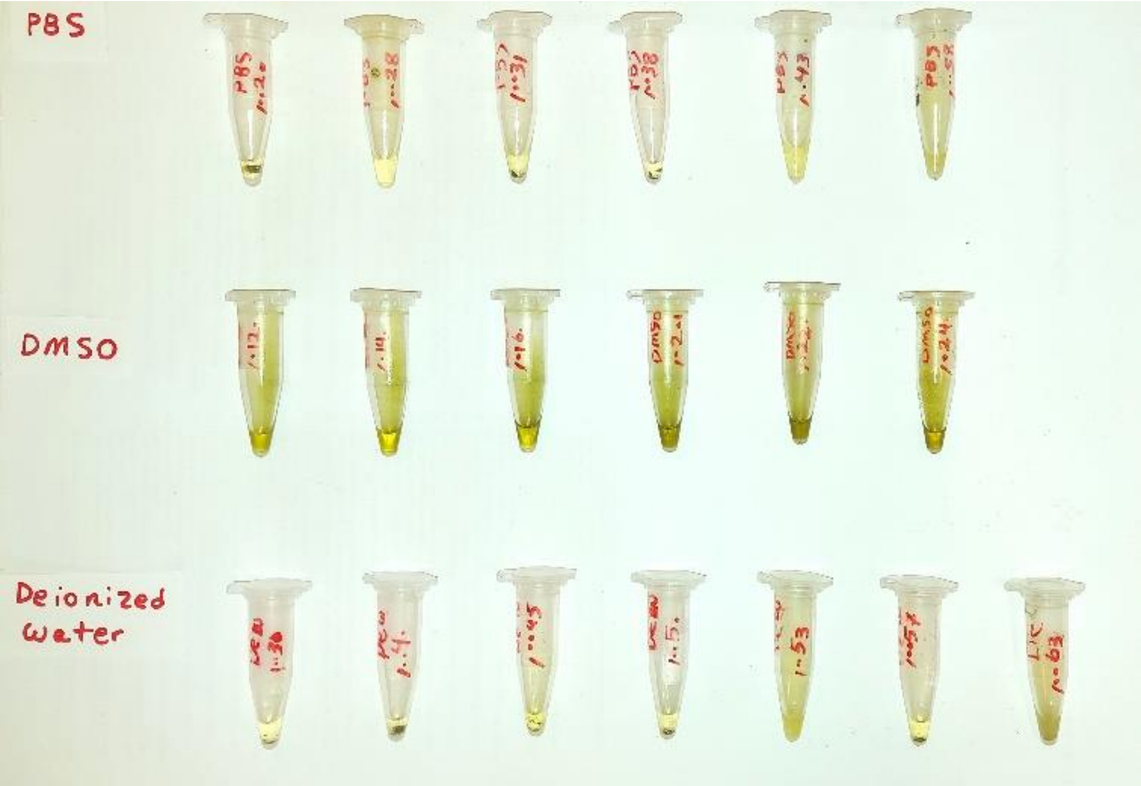


**Figure. S1** Determining the suitable solvent for dissolving the dry extract. The extracts were dissolved at various weights in 200 µL of the three solvents

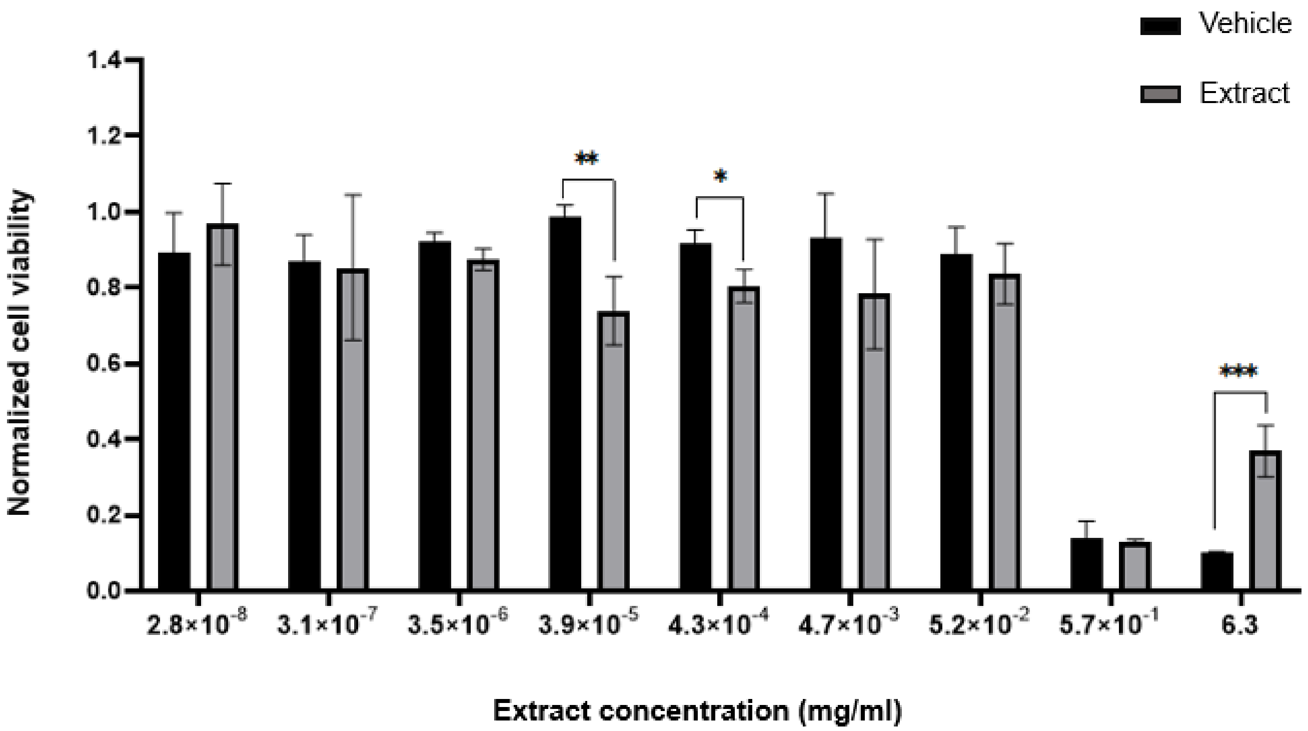


**Figure. S2** The effect of *C. sativa* seed extract dissolved in DMSO on the viability of PANC-1 cells. Quantitative assessment of viability of PANC-1 cells after treatment with the extract dissolved in DMSO for 72h
